# Regulation of blood–brain barrier integrity by microbiome-associated methylamines and cognition by trimethylamine *N*-oxide

**DOI:** 10.1101/2021.01.28.428430

**Authors:** Lesley Hoyles, Matthew G. Pontifex, Ildefonso Rodriguez-Ramiro, M. Areeb Anis-Alavi, Khadija S. Jelane, Tom Snelling, Egle Solito, Sonia Fonseca, Ana L. Carvalho, Simon R. Carding, Michael Müller, Robert C. Glen, David Vauzour, Simon McArthur

## Abstract

**Background:** Communication between the gut microbiota and the brain is primarily mediated *via* soluble microbe-derived metabolites, but the details of this pathway remain poorly defined. Methylamines produced by microbial metabolism of dietary choline and L-carnitine have received attention due to their proposed association with vascular disease, but their effects upon the cerebrovascular circulation have hitherto not been studied.

**Results:** Here we use an integrated *in vitro*/*in vivo* approach to show that physiologically relevant concentrations of the dietary methylamine trimethylamine *N*-oxide (TMAO) enhanced blood-brain barrier (BBB) integrity and protected it from inflammatory insult, acting through the tight junction regulator annexin A1. In contrast, the TMAO precursor trimethylamine (TMA) impaired BBB function and disrupted tight junction integrity. Moreover, we show that long-term exposure to TMAO protects murine cognitive function from inflammatory challenge, acting to limit astrocyte and microglial reactivity in a brain region-specific manner.

**Conclusion:** Our findings demonstrate the mechanisms through which microbiome-associated methylamines directly interact with the mammalian BBB, with consequences for cerebrovascular and cognitive function.

## INTRODUCTION

As the role of the gut microbiota in host physiology and disease is categorised, novel pathways through which these interactions are mediated continue to emerge. We and others recently identified the blood–brain barrier (BBB) as a target for gut microbe-derived short-chain fatty acid (SCFA) activity, with butyrate and propionate acting to promote BBB integrity and protect the cerebral vasculature from insult [1,2]. SCFAs represent just one of many classes of gut microbe-derived metabolites, with little known as to how these other classes may influence BBB function.

Dietary methylamines, such as choline, phosphatidylcholine, betaine and trimethylamine-*N*-oxide (TMAO), are a class of metabolites receiving considerable attention as modulators of vascular function [3,4], although the mechanism(s) by which they affect human physiology remain poorly understood. The aforementioned methylamines can be broken down by members of the gut microbiota into trimethylamine (TMA) [5], which is carried from the gut through the portal vasculature to the liver and rapidly converted into TMAO by flavin monooxygenases [6]. TMAO then enters the systemic circulation, reaching fasting plasma concentrations of between 2 and 40 µM in humans [7–9], prior to excretion through the urine [5]. Approximately ten-fold lower concentrations of TMA compared with TMAO are found in the circulation under normal physiological conditions.

Early observational work reported an association between atherosclerosis and elevated levels of TMAO [10,11]. Similarly, pre-clinical studies demonstrate the damaging effects of supraphysiological TMAO doses in atherosclerosis-prone mice [12] and upon thrombus formation [13]. Despite this, the impact of TMAO upon the vasculature remains uncertain, with a number of detailed studies encompassing both human and murine systems having failed to replicate these initial findings [14], instead suggesting that this negative relationship disappears upon correction for renal function [4,15–17] and thus indicating that raised TMAO levels may in fact reflect impaired excretion rather than being a causative factor in disease. Moreover, protective roles for TMAO have been reported in rodent models of hypertension [18], atherosclerosis [19] and non-alcoholic steatohepatitis [20] and we have previously shown TMAO to improve glucose homeostasis and insulin secretion in mice fed a high-fat diet [21]. Perhaps helping to clarify this apparent contradiction, recent studies have established that intravenous treatment of rats with the TMAO precursor TMA, but not TMAO itself, increases mean arterial blood pressure [22]. Notably, the majority of reports describing associations of plasma TMAO with cardiovascular disease have not concurrently monitored levels of TMA; TMA but not TMAO has been shown to associate with severe aortic stenosis [22] and gestational diabetes risk [23].

Beyond vascular health, dietary methylamines have implications for cognition, with a positive correlation observed between choline intake and cognitive function in both humans [24,25] and mice [26,27]. In contrast, cerebrospinal fluid TMAO levels have been indicated as predictive of cognitive decline in Alzheimer’s disease [28], while suppression of microbial TMA/TMAO production improves cognitive function in the murine APP/PS1 model of Alzheimer’s disease [29]. Given the disparities in the literature regarding the effects of methylamines upon the vasculature, and our increasing awareness of the BBB as a major actor in the pathology of multiple neurological conditions, we investigated the effects of physiologically relevant concentrations of TMAO and its precursor TMA upon BBB integrity and cognitive behaviour.

## METHODS

### Endothelial cell culture

The human cerebromicrovascular endothelial cell line hCMEC/D3 was maintained and treated as described previously [2,30]. Cells bearing shRNA sequences targeting annexin A1 (*ANXA1*) or non-specific scramble sequences were produced as described previously [31]; the degree of *ANXA1* knock-down was confirmed by flow cytometry analysis (Suppl. Fig. 1). For all lines, cells were cultured to confluency in complete EBM-2MV microvascular endothelial cell growth medium (Promocell GmbH, Heidelberg, Germany), whereupon medium was replaced by EBM-2MV without VEGF and cells were further cultured for a minimum of 4 days to enable intercellular tight junction formation prior to experimentation.

### In vitro barrier function assessments

Paracellular permeability and transendothelial electrical resistance (TEER) were measured on 100 % confluent hCMEC/D3 cultures polarised by growth on 24-well plate polyethylene terephthalate (PET) transwell inserts (surface area: 0.33 cm^2^, pore size: 0.4 μm; Greiner Bio-One GmbH, Kremsmünster, Austria) coated with calf-skin collagen and fibronectin (Sigma-Aldrich, UK). The permeability of hCMEC/D3 cell monolayers to 70 kDa FITC-dextran (2 mg/ml) was measured as described previously [31–33]. TEER measurements were performed using a Millicell ERS-2 Voltohmmeter (Millipore, Watford, UK) and were expressed as Ω.cm^2^. In all cases, values obtained from cell-free inserts similarly coated with collagen and fibronectin were subtracted from the total values. In some cases, barrier integrity was tested by challenge with bacterial lipopolysaccharide (LPS). Confluent hCMEC/D3 monolayers were treated with TMAO or TMA for 12 h, whereupon LPS (*Escherichia coli* O111:B4; 50 ng/ml, comparable to circulating levels of LPS in human endotoxemia [34]) was added for a further 12 h, without wash-out. Barrier function characteristics were then interrogated as described above.

### Cell adhesion assays

hCMEC/D3 cells were cultured to confluency on transwell inserts (0.4 µm pore size, 0.33 cm^2^ diameter, Greiner Bio-One Gmbh, Austria) prior to 16 h treatment with 10 ng/ml TNFα. Monolayers were then incubated for 2 h with U937 monocytic cells pre-labelled according to manufacturer’s instructions with CMFDA cell tracker dye (ThermoFisher Scientific, UK). Co-cultures were washed vigorously with ice-cold PBS three times and fixed by incubation for 10 min in 1 % formaldehyde in 0.1 M PBS. Co-cultures were mounted and examined using an Axiovert 200M inverted microscope (Zeiss) equipped with a 20x objective lens. Images were captured with ZEN imaging software (Carl Zeiss Ltd, UK) and analysed using ImageJ 1.53c (National Institutes of Health, USA).

### Microarrays

hCMEC/D3 cells were grown on 6-well plates coated with calf-skin collagen (Sigma-Aldrich, Gillingham, UK), and collected in TRIzol (Thermo-Fisher Scientific, UK) as described previously [2]. Total RNA was extracted using a TRIzol Plus RNA purification kit (Thermo-Fisher Scientific, UK) and quantified using a CLARIOstar spectrophotometer equipped with an LVis microplate (BMG Labtech GmbH, Germany).

Hybridization experiments were performed by Macrogen Inc. (Seoul, Republic of Korea) using Illumina HumanHT-12 v4.0 Expression BeadChips (Illumina Inc., San Diego, CA) as described previously [2].

### Processing and analyses of array data

Raw data supplied by Macrogen were quality-checked, log_2_-transformed and loess-normalized (2 iterations) using affy [35]. Probe filtering and matching of probes not classified as ‘Bad’ or ‘No match’ to Entrez identifiers were done as described previously [2]. Average gene expression values were used for identification of differentially expressed genes. Array data have been deposited in ArrayExpress under accession number E-MTAB-6662. Normalized data are available (Supplementary Table 1).

Enrichr [36,37] was used to perform Gene Ontology (GO) analysis. Signaling Pathway Impact Analysis (SPIA) was used to determine whether Kyoto Encyclopedia of Genes and Genomes (KEGG) pathways were activated or inhibited in hCMEC/D3 cells exposed to TMAO or TMA [38]. Human KEGG pathways (KGML format) downloaded from the KEGG PATHWAY database [39] were used for network (KEGGgraph, RBGL [40]) analysis.

### Immunofluorescence microscopy

hCMEC/D3 cells were cultured on 24-well plate PET transwell inserts (surface area: 0.33 cm^2^, pore size: 0.4 μm; Greiner Bio-One GmbH, Kremsmünster, Austria) coated with calf-skin collagen and fibronectin (Sigma-Aldrich, UK), prior to immunostaining according to standard protocols [2,31] and using a primary antibody directed against zonula occludens-1 (ZO-1; 1:100, ThermoFisher Scientific, UK) or Alexafluor 488-conjugated phalloidin (1:140, ThermoFisher Scientific, UK). Nuclei were counterstained with DAPI (Sigma-Aldrich, UK). Images were captured using an LSM880 confocal laser scanning microscope (Carl Zeiss Ltd, Cambridge, UK) fitted with 405 nm and 488 nm lasers, and a 63x oil immersion objective lens (NA, 1.4 mm, working distance, 0.17 mm). Images were captured with ZEN imaging software (Carl Zeiss Ltd, UK) and analysed using ImageJ 1.53c (National Institutes of Health, USA).

### Flow cytometry analysis

Following experimental treatment, hCMEC/D3 cells were detached using 0.05 % trypsin and incubated with an unconjugated rabbit polyclonal antibody directed against ANXA1 (1:1000, ThermoFisher Scientific, UK) on ice for 30 min, followed by incubation with an AF488-conjugated goat anti-rabbit secondary antibody (1:500, ThermoFisher Scientific, UK). Similarly detached hCMEC/D3 cells were incubated with APC-conjugated mouse monoclonal anti-BCRP (1:100, BD Biosciences, Oxford, UK), or PE-conjugated mouse monoclonal anti-MDR1A (1:100, BD Biosciences, UK) antibodies on ice for 30 min, alongside fluorescence minus one controls. Immunofluorescence was analysed for 20,000 events per treatment using a BD FACSCanto II (BD Biosciences, UK) flow cytometer; data were analysed using FlowJo 8.0 software (Treestar Inc., CA, USA).

### Efflux transporter assays

Activity of the major efflux transporters P-glycoprotein and Breast Cancer Resistance Protein (BCRP) was determined through the use of commercially available assays (PREDEASY™ ATPase Assay Kits, Solvo Biotechnology Inc., Budapest, Hungary), performed according to the manufacturer’s instructions. Stepwise dose–response curves centred around reported physiological circulating concentrations of TMA (4.9 nM – 10.8 µM) and TMAO (0.5 µM – 1.08 mM) were constructed (*n* = 4) to investigate inhibitory effects of the methylamines upon transporter activity.

### ELISA

Culture medium ANXA1 content was assayed by specific ELISA as described previously [41]. Serum TNFα and IL-1β concentrations were assayed using commercial ELISA kits according to the manufacturer’s instructions (ThermoFisher Scientific, UK).

### Animal experiments

All animal experiments were performed according to the UK Animals (Scientific Procedures) Act of 1986, under UK Home Office Project Licences PFA5C4F4F (short term studies) and 70/8710 (long term studies), following ethical review by the Animal Welfare and Ethical Review Boards of Queen Mary, University of London or the University of East Anglia, respectively. Wild-type male C57Bl/6J mice (Charles River Ltd, Harlow, UK) aged 8 weeks at the start of procedures were used throughout, with a group size of n=5-6 for short term studies and n=8 for long-term/behavioural analyses. Animals were housed in individually ventilated cages on a daily 12 h:12 h light/dark cycle with, unless otherwise indicated, *ad libitum* access to standard mouse chow and drinking water. Experimental procedures were started at 9 am to minimise variation associated with circadian rhythms.

### Assessment of acute effects of TMAO on BBB integrity

Mice (n=5-6 per group) were injected intraperitoneally (i.p.) with 1.8 mg/kg body weight TMAO in 100 µl saline vehicle, a dose calculated to approximate human circulating TMAO levels [42], followed 2 h, 6 h or 24 h later by assessment of Evans blue extravasation as described below. Alternatively, mice were injected i.p. with 3 mg/kg body weight LPS or 100 µl 0.9% saline vehicle, followed 2 h later by i.p. injection of either 1.8 mg/kg body weight TMAO or 100 µl 0.9% saline vehicle for assessment of Evans blue extravasation 2 h later. In both experiments, one hour before assessment animals were injected i.p. with 100 µl of a 2 % (w/v) solution of Evans blue dye in 0.9 % saline (Sigma–Aldrich Ltd, Poole, UK). Dye was permitted to circulate for 1 h before animals were transcardially perfused with 0.9 % saline at 4 °C to remove circulating dye. Brains were removed, bisected and homogenized by maceration in 0.9 % saline. Suspended macromolecules were precipitated by incubation with 60 % trichloroacetic acid, and dye content of resulting supernatants was detected using a CLARIOstar spectrophotometer (BMG Labtech GmbH, Germany) alongside a standard curve of defined concentrations of Evans blue in the same buffer. Brain Evans blue content was expressed as µg of dye per mg of brain tissue, normalized to circulating plasma concentrations.

### Long-term LPS and TMAO treatments

To assess the long-term impact of both LPS and TMAO on cognitive performance, mice were divided into four groups (n=8 per group): 1) Water + PBS; 2) Water + TMAO; 3) LPS + PBS; 4) LPS + TMAO. C57Bl/6 mice were administered phosphate-buffered saline (PBS) or LPS (*Escherichia coli* O55:B5, Sigma-Aldrich, UK) via i.p. injection (0.5 mg/kg/wk) for 8 weeks [43]. A final LPS treatment was administered the day before sacrifice for nine total injections. Body weights were recorded prior to each injection. Starting on the day of the first saline/LPS injection, TMAO was provided in the drinking water (500 mg/L), with water bottles being replaced every other day. Drinking volumes were recorded before bottle change.

### Processing and analyses of RNAseq data

Mice were transcardially perfused with 0.9 % saline at 4 °C to remove circulating blood, and brains were removed and collected into RNAlater (Thermofisher Scientific Ltd., UK) prior to storage at −20 °C for later analysis. Whole brain total RNA was extracted using a PureLink RNA Mini Kit (Thermofisher Scientific Ltd., UK) and quantified using a CLARIOstar spectrophotometer equipped with an LVis microplate (BMG Labtech GmbH, Germany). RNA samples (*n*=3 TMAO, *n*=3 control) were sent to Macrogen Inc. (Republic of Korea) where they were subject to quality checks (RIN analysis); libraries were prepared (TruSeq Stranded mRNA LT Sample Prep Kit) for paired-end (2x 100 nt) sequencing on an Illumina HiSeq 4000 apparatus. Raw RNAseq sequence data (delivered in fastq format) were processed in house as follows. Reads were mapped onto the mouse genome (mm10) using HISAT2 v2.1.0 [44]. Number of reads in each sample that mapped to genes in the BAM files returned by HISAT2 was determined using featureCounts v1.6.4 [45]. Entrez gene identifiers were converted to gene symbols using *Mus musculus* annotations downloaded from NCBI on 26 November 2020; only those genes with valid Entrez gene identifiers were retained in analyses. Raw RNAseq data have been deposited with ArrayExpress under accession number E-MTAB-9869. Significantly differentially expressed genes (*P*<0.1) were analysed by mouse KEGG pathway over-representation analysis using Enrichr and manual curation.

### Behavioural analyses

Behavioural tests were performed in the order they are introduced below. Apparatus was cleaned using 70 % ethanol upon completion of each trial, eliminating any residual odour.

Open field test (OFT) was conducted as previously described [46]. Briefly, mice were placed in the centre of the OFT, a grey 50 x 50 x 50 cm apparatus illuminated with low lux (100 lux) lighting. Total travel distance and time spent in the centre of the field was determined at 5 min with a video tracking system (Smart 3.0 tracking software, Panlab, Kent, UK).

The novel object recognition (NOR), a measure of recognition memory, was performed as described previously [47,48], with slight modifications. Briefly, on day 1 mice were habituated in a grey 50 x 50 x 50 cm apparatus illuminated with low lux (100 lux) lighting, mice were placed into the empty maze and allowed to move freely for 10 min. On day 2, mice were conditioned to a single object for a 10 min period. On day 3, mice were placed into the same experimental area in the presence of two identical objects for 15 min, after which they were returned to their respective cages and an inter-trial interval of 1 h was observed. One familiar object was replaced with a novel object, with the position of the novel object (left or right) being randomized between each mouse and group tested. Mice were placed back within the testing area for a final 10 min. Videos were analysed for a 5 min period, after which if an accumulative total of 15 s with both objects failed to be reached, analysis continued for the full 10 min or until 15 s was achieved. Those not achieving 15 s were excluded from the analysis [49]. A discrimination index (DI) was calculated as follows: DI = (TN−TF)/(TN+TF), where TN is the time spent exploring the novel object and TF is the time spent exploring the familiar object.

Y-maze spontaneous alternation test, a measure of spatial working memory, was performed on the final day of behavioural testing as previously described [50]. Briefly, the Y-maze apparatus comprised white Plexiglas (dimensions 38.5 × 8 × 13 cm, spaced 120° apart) and was illuminated with low lux (100 lux) lighting. Mice were placed in the maze and allowed to explore freely for 7 min while tracking software recorded zone transitioning and locomotor activity (Smart 3.0 tracking software, Panlab, Kent, UK). Spontaneous alternation was calculated using the following formula: Spontaneous Alternation = (Number of alternations/ Total Arm entries - 2) x 100.

### Extravasation assay and sample processing following long-term treatment

Twenty-four hours after the final injection of LPS, mice were injected i.p. with 200 µl of 2 % sodium fluorescein in sterile ddH_2_O and anesthetized 30 min later with isoflurane (1.5 %) in a mixture of nitrous oxide (70 %), and oxygen (30 %). Once sedated, blood was collected by cardiac puncture and centrifuged at 1,500 *g* for 15 min at 4 °C to collect the serum. The samples were analysed immediately for sodium fluorescein extravasation or snap-frozen in liquid nitrogen and stored at −80 °C until further analysis.

Mice were then transcardially perfused with saline containing 10 kU/ml heparin (Sigma, Devon, UK). Dissected left hemi-brains were fixed in 4% PFA for 24 h and embedded into paraffin before being processed for immunohistochemical analysis. Right hemi-brains were stored at −80 °C until further analysis; cerebellums were processed immediately for the sodium fluorescein extravasation assay. Cleared volume of sodium fluorescein that passed from the plasma into the brain was calculated as described previously [43].

### Ex vivo immunohistochemical analysis

Paraffin-embedded brains were sectioned (5 µm) using a rotary microtome and collected onto glass microscope slides. Following deparaffinisation using xylene and rehydration using graded ethanol:water solutions, heat-mediated antigen retrieval was performed by incubation in 10 mM Tris base, 1 mM EDTA, 0.05 % Tween-20, pH 9.0 at 90 °C for 20 min. Once cooled, endogenous peroxide activity was quenched by incubation for 15 min in 0.3 % H_2_O_2_ in Tris-buffered saline (TBS; 50 mM Tris base, 150 mM NaCl, pH 7.4). Sections were permeabilised and blocked by incubation in TBS containing 0.025% triton X-100 and 10 % normal goat serum for 30 min, prior to overnight treatment at 4 °C with rabbit anti-murine primary antibodies raised against GFAP (1:1000, ab7260, Abcam Ltd, UK) or Iba1 (1:1000, 019-19741, FUJIFILM Wako Pure Chemical Corporation, Japan) diluted in TBS containing 1 % normal goat serum, 0.025 % Triton X-100, pH 7.4. Sections were washed thoroughly with TBS containing 1 % normal goat serum and incubated for 1 h at room temperature with a horseradish peroxidase-conjugated goat anti-rabbit antibody (1:500, Stratech Scientific, UK) diluted in TBS containing 1 % normal goat serum, 0.025 % Triton X-100, pH 7.4). Sections were thoroughly washed in TBS and peroxidase staining was developed using diaminobenzidine hydrochloride and H_2_O_2_. Sections were dehydrated with graded ethanol:water solutions, cleared with xylene and mounted under DPX for microscopic examination. Brightfield images were captured using a using a Nikon Eclipse 80i Stereology Microscope fitted with an Optronics Camera, using a 20x objective, and analysed with ImageJ 1.53 k software (National Institutes of Health, USA).

### Statistical analyses

Sample sizes were calculated to detect differences of 15 % or more with a power of 0.85 and α set at 5 %, calculations being informed by previously published data [2,31]. *In vitro* experimental data (except those for *in vitro* microarray experiments) are expressed as mean ± SEM, with a minimum of *n* = 3 independent experiments performed in triplicate for all studies. In all cases, normality of distribution was established using the Shapiro–Wilks test, followed by analysis with two-tailed Student’s *t*-tests to compare two groups or, for multiple comparison analysis, 1- or 2-way ANOVA followed by Tukey’s HSD *post hoc* test, or Dunnett’s test for dose-response experiments. Where data were not normally distributed, non-parametric analysis was performed using the Wilcoxon signed rank test. A *P* value of less than or equal to 5 % was considered significant. Differentially expressed genes were identified in microarray data using LIMMA [51]; *P* values were corrected for multiple testing using the Benjamini– Hochberg procedure (False Discovery Rate); a *P* value of less than or equal to 10 % was considered significant in this case; *n* = 5 for control, TMAO and TMA groups. Significantly differentially expressed genes (P_FDR_<0.1) in RNAseq data (Supplementary Table 11) were identified using DESeq2 v1.22.1 [52].

## RESULTS

To provide an initial assessment of the effects of the methylamines TMA and TMAO upon the BBB we used a well-established *in vitro* BBB model, hCMEC/D3 immortalised human cerebromicrovascular cell monolayers grown under polarising conditions on a Transwell filter, examining two key barrier properties: paracellular permeability to a protein-sized tracer and TEER. Exposure of hCMEC/D3 cells for 24 h to TMA (0-40 µM) caused a clear dose-dependent increase in paracellular permeability to 70 kDa FITC-dextran (Fig. 1A), with normal circulating levels (0.4 µM) of TMA and upwards significantly enhancing permeability. In contrast, exposure for 24 h to TMAO (0-4000 µM) caused a biphasic dose-dependent response (Fig. 1A), with normal circulating concentrations (4-40 µM) significantly reducing permeability to the tracer, an effect lost at 2.5-fold greater TMAO concentrations and reversed at 100-fold greater TMAO (4 mM), where a significant increase in paracellular permeability was apparent. In contrast, TMA had no effect upon TEER at any concentration studied, while TMAO enhanced TEER by approximately 65%, an effect that was notably dose-independent (Fig. 1B).

**Fig. 1.**
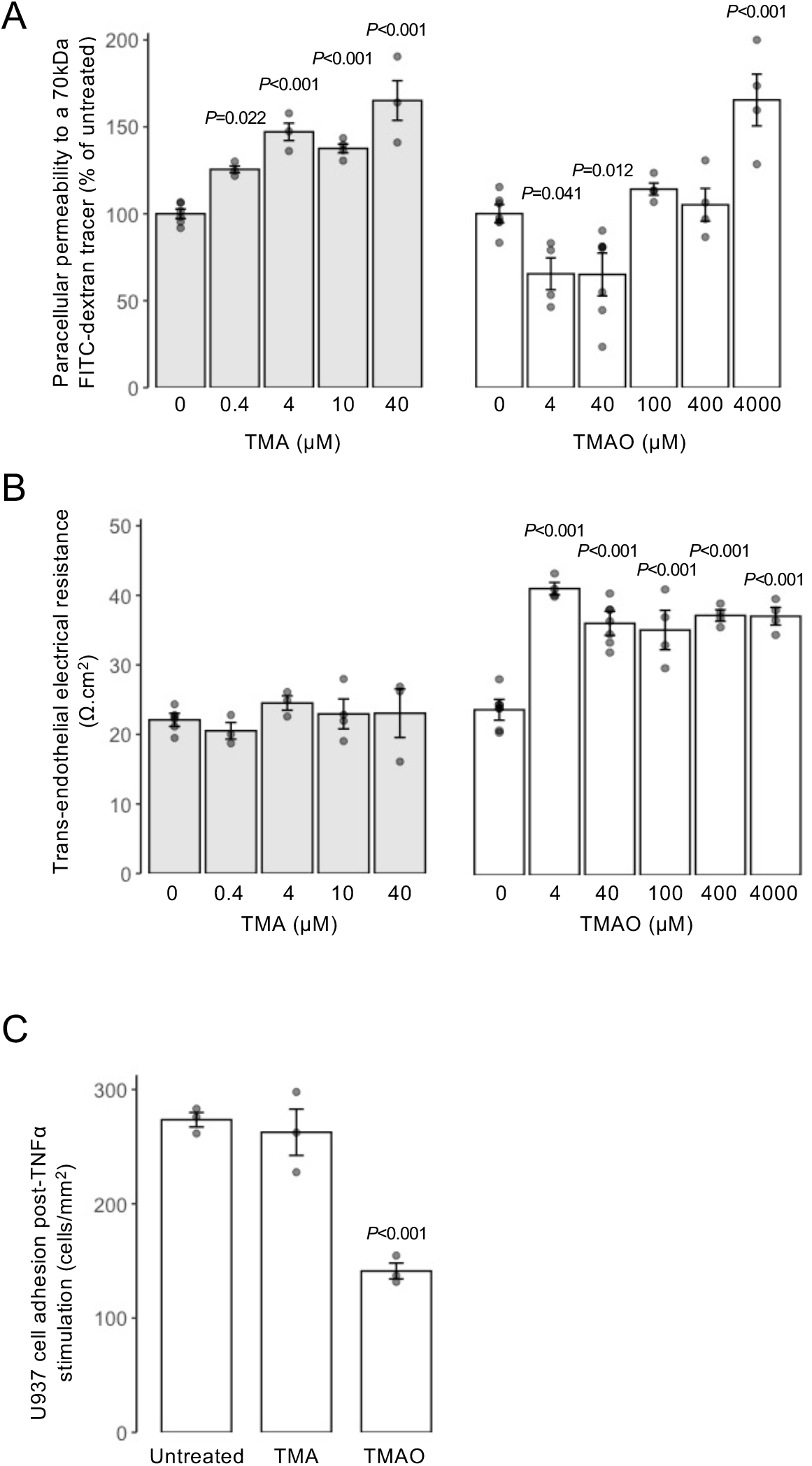
Effects of TMAO and TMA on integrity of hCMEC/D3 cell monolayers. (A) Assessment of paracellular permeability of hCMEC/D3 monolayers to a 70 kDa FITC-dextran tracer following treatment for 24 h with varying doses of TMA (0.4 – 40 µM) or TMAO (4 – 4000 µM). Data are expressed as mean ± s.e.m., *n*=4 independent experiments. (B) Assessment of TEER of hCMEC/D3 monolayers to a 70kDa FITC-dextran tracer following treatment for 24 h with varying doses of TMA (0.4 – 40 µM) or TMAO (4 – 4000 µM). Data are expressed as mean ± s.e.m., *n*=4 independent experiments. (C) Adhesion of U937 monocytic cells to TNFα-stimulated hCMEC/D3 monolayers (10 ng/ml, 16 h) that had been treated or not for 24 h with 0.4 µM TMA or 40 µM TMAO. Data are expressed as mean ± s.e.m., *n*=3 independent experiments.

The physical barrier that the BBB provides is only one aspect by which it separates the brain parenchymal environment from the periphery, equally important is the immunological barrier that it represents. To model this, we employed a simple system in which adhesion of CMFDA-labelled U937 monocytic cells to TNFα-activated (10 ng/ml, 16 h) hCMEC/D3 monolayers was quantified in response to TMA or TMAO treatment. Treatment with a physiologically relevant concentration of TMA (0.4 µM [42], 24 h post-TNFα) had no effect on the density of adherent U937 cells, but exposure of hCMEC/D3 monolayers to physiological levels of TMAO (40 µM [42], 24 h post-TNFα) significantly reduced U937 cell adhesion by approximately 50 % compared to cultures stimulated with TNFαalone (Fig. 1C).

The endothelial cells of the BBB express numerous efflux transporter proteins that serve to limit entry of endogenous and exogenous molecules into the parenchyma, with BCRP and P-glycoprotein being two of the most important. Consequently, we examined whether treatment with TMA or TMAO affected function or expression of either of these two transporters. Using commercially available *in vitro* assays, neither methylamine affected BCRP nor P-glycoprotein activity across a wide concentration range (TMA: 4.9 nM to 10.8 µM; TMAO 0.5 µM to 1.08 mM) (Suppl. Fig. 2A-D). Similarly, treatment of hCMEC/D3 cells for 24 h with physiologically relevant concentrations of TMA (0.4 µM) or TMAO (40 µM) was without effect on cell surface expression of either BCRP or P-glycoprotein (Suppl. Fig. 2E-F).

### Methylamine-induced changes in gene expression

Having identified significant TMA-/TMAO-induced functional changes in endothelial barrier characteristics *in vitro*, we undertook a microarray analysis of hCMEC/D3 cells treated with either TMA (0.4 µM, 24 h) or TMAO (40 µM, 24 h) to investigate the transcriptional changes underlying these effects. Treatment with TMA had a significant (P_FDR_<0.1) effect on 49 genes, with the expression of 39 upregulated and 10 downregulated (Fig 2A, Supplementary Table 2). In contrast, treatment with TMAO had a significant (P_FDR_<0.1) effect on 440 genes with 341 upregulated and 99 downregulated (Fig. 2B, Supplementary Table 3). *FMO3* gene expression was not affected by TMA or TMAO at the physiological concentrations employed (Suppl. Fig. 3).

**Fig. 2.**
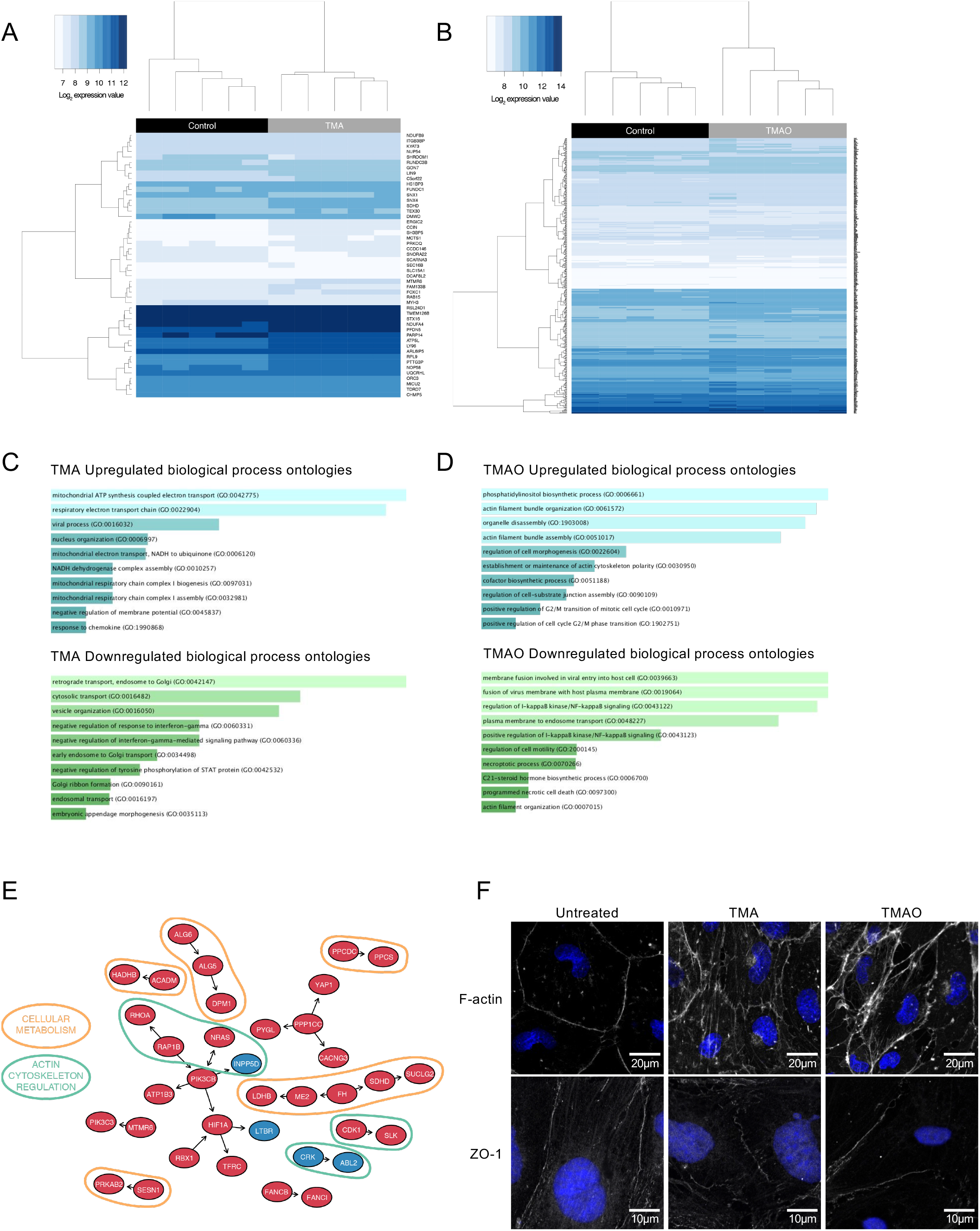
Effects of TMA and TMAO on gene expression in hCMEC/D3 cells. (A) Heatmap showing expression of the 49 genes found to be significantly (P_FDR_<0.1) differentially expressed upon exposure of hCMEC/D3 cells to 0.4 µM TMA (*n*=5 per group). (B) Heatmap showing expression of the 440 genes found to be significantly (P_FDR_<0.1) differentially expressed upon exposure of hCMEC/D3 cells to 40 µM TMAO (*n*=5 per group). (C) Biological processes associated with genes found to be significantly upregulated (*n*=39) or downregulated (*n*=10) upon exposure of cells to TMA. (D) Biological processes of genes found to be significantly upregulated (*n*=341) or downregulated (*n*=99) upon exposure of cells to TMAO. Images in (C, D) shown based on Enrichr *P* value ranking from GO analysis. (E) Topological analysis of the KEGG networks associated with the 440 genes whose expression was significantly affected upon exposure of cells to TMAO (blue, significantly downregulated; red, significantly upregulated); genes of similar cellullar role are highlighted. (F) Confocal microscopic analysis of expression of fibrillar actin (F-actin) and the tight junction component zonula occludens-1 (ZO-1) in hCMEC/D3 cells following treatment for 24 h with 0.4 μM TMA or 40 µM TMAO. Images are representative of at least three independent experiments.

SPIA of the 440 TMAO-affected genes showed activation of the tight junction pathway (P = 0.031), but significance was lost after correction for multiple testing (Supplementary Table 4). No pathways were shown to be activated or inactivated by the 49 TMA-affected genes (data not shown).

Gene ontology (GO) analysis was performed on TMA- and TMAO-regulated genes using Enrichr [36,37]. TMA up-regulated and down-regulated genes were significantly (P_FDR_<0.2) associated with processes indicative of a degree of cellular stress (Fig 2C, Supplementary Table 5, Supplementary Table 6). In contrast, genes up-regulated by TMAO treatment were associated with regulation of the cytoskeleton and cell morphology and with actin bundle formation (P_FDR_<0.2), whereas pathways associated with inflammatory signalling were down-regulated (Fig 2D, Supplementary Table 7, Supplementary Table 8).

We then assessed the topology of a directional network of the 440 TMAO-associated genes mapped onto all human KEGG pathways. In line with the GO analysis described above, a number of genes of differing function were regulated by TMAO treatment, with two principal groupings being particularly evident, namely those associated with aspects of cellular metabolism and with regulation of actin cytoskeletal dynamics (Figure 2E). Finally, we compared the 19,309 genes represented on the microarray with a set of 203 genes [2] known to be associated with the BBB. While TMA treatment had no significant effects on expression of these genes (Supplementary Table 9), TMAO significantly (P_FDR_<0.1) upregulated expression of four genes from this set associated with transporter proteins and barrier integrity (Table 1, Supplementary Table 10).

**Table 1.** BBB-associated genes whose expression was upregulated upon exposure of hCMEC/D3 cells to TMAO.

| Gene | Entrez ID | Description | Log<sub>2</sub> fold change | Category | P<sub>FDR</sub> |
| --- | --- | --- | --- | --- | --- |
| *TFRC* | 7037 | Transferrin receptor | 0.23 | Transporter proteins | 0.054 |
| *ABCC4* | 10257 | ATP binding cassette subfamily C member 4 | 0.20 | Transporter proteins | 0.088 |
| *ANXA1* | 301 | Annexin A1 | 0.16 | Cell Adhesion/Junctional proteins/Cytoskeletal factors | 0.088 |
| *CDH2* | 1000 | Cadherin 2 | 0.31 | Cell Adhesion/Junctional proteins/Cytoskeletal factors | 0.095 |

Given these transcriptional indications, and the fact that the restrictive properties of the BBB are largely governed by inter-endothelial cell tight junctions linked *via* the zonula occludens complex to the actin cytoskeleton [53], we hypothesised that TMA and TMAO may affect barrier permeability through modification of the links between tight junctions and the actin cytoskeleton. Confocal immunofluorescence microscopy of hCMEC/D3 monolayers treated with a physiologically relevant concentration of TMA (0.4 µM, 24 h) or TMAO (40 µM, 24 h) revealed clear changes to both ZO-1 and fibrillar actin disposition within cells (Fig. 2F). Compared to untreated cells in which both ZO-1 and F-actin fibres clearly defined the cellular perimeter, cells treated with TMA exhibited a broken, patchy distribution of perimeter ZO-1 expression, and the appearance of marked cytoplasmic F-actin stress fibres. In contrast, cells treated with TMAO showed little change in ZO-1 distribution, but a marked enhancement of cortical F-actin fibre thickness and intensity.

### The actions of TMAO are mediated through annexin A1 signalling

Of the four BBB-associated genes identified as upregulated by TMAO, *ANXA1* is of particular interest as we have previously shown this protein to regulate BBB tightness *in vitro* and *in vivo* through modulation of the actin cytoskeleton [54]. Examination of ANXA1 expression in hCMEC/D3 cells revealed that while total cellular levels of the protein were not changed by either TMA (0.4 µM, 24 h) or TMAO (40 µM, 24 h) treatment (Fig. 3A), TMA significantly suppressed and TMAO significantly augmented medium ANXA1 content (Fig. 3B), a finding of interest given that autocrine/paracrine effects are a major route of ANXA1 action [55].

**Fig. 3.**
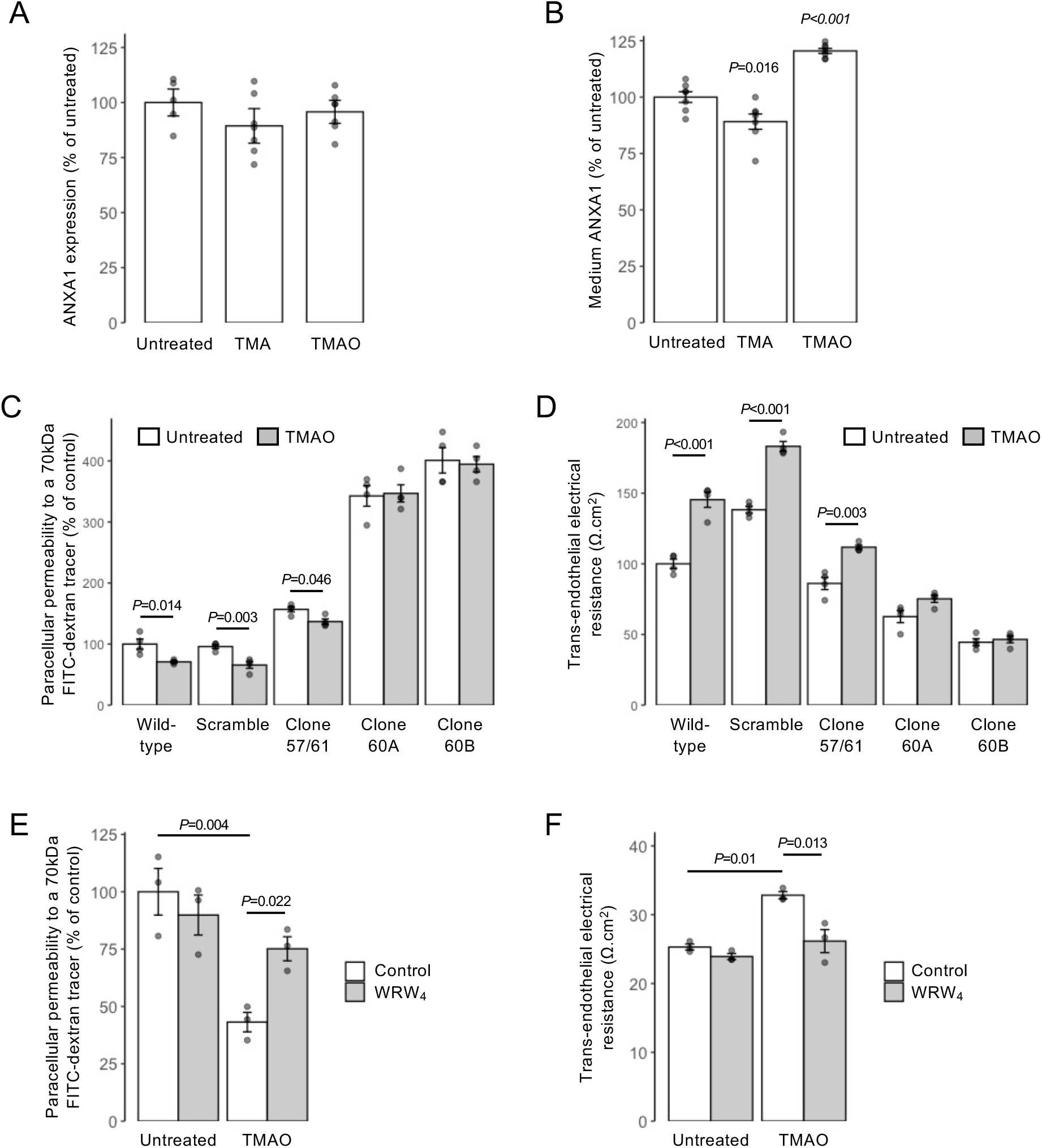
Annexin A1 (ANXA1) signalling mediates effects of TMAO on hCMEC/D3 cells. (A) Total cellular expression of ANXA1 in hCMEC/D3 cells treated for 24 h with 0.4 µM TMA or 40 µM TMAO. Data are expressed as mean ± s.e.m., *n*=5-7 independent experiments. (B) Medium ANXA1 content of hCMEC/D3 monolayers treated for 24 h with 0.4 µM TMA or 40 µM TMAO. Data are expressed as mean ± s.e.m., *n*=7 independent experiments. (C) Assessment of paracellular permeability of monolayers of wild-type hCMEC/D3 cells, or hCMEC/D3 cells stably transfected with either a scramble shRNA sequence, or one of three shRNA sequences targeting ANXA1 (clone 57/61 – 20.6 ± 5.6% reduction, clone 60A – 47.3 ± 1.5% reduction, clone 60B – 67.5 ± 1.1% reduction) to a 70kDa FITC-dextran tracer following treatment for 24 h with 40 µM TMAO. Data are expressed as mean ± s.e.m., *n*=4 independent experiments. (D) Assessment of TEER of monolayers of wild-type hCMEC/D3 cells, or hCMEC/D3 cells stably transfected with either a scramble shRNA sequence, or one of three shRNA sequences targeting ANXA1 (clone 57/61 – 20.6 ± 5.6% reduction, clone 60A – 47.3 ± 1.5% reduction, clone 60B – 67.5 ± 1.1% reduction) following treatment for 24 h with 40 µM TMAO. Data are expressed as mean ± s.e.m., *n*=4 independent experiments. (E) Assessment of paracellular permeability of hCMEC/D3 cells to a 70kDa FITC-dextran tracer following treatment for 24 h with 40 µM TMAO, with or without 10 min pre-treatment with the FPR2 antagonist WRW_4_ (10 µM). Data are expressed as mean ± s.e.m., *n*=3 independent experiments. (F) Assessment of TEER of hCMEC/D3 cells following treatment for 24 h with 40 µM TMAO, with or without 10 min pre-treatment with the FPR2 antagonist WRW_4_ (10 µM). Data are expressed as mean ± s.e.m., *n*=3 independent experiments.

To establish the importance of ANXA1 in mediating the effects of TMAO, we investigated the effects of its depletion through use of hCMEC/D3 clones stably transfected with shRNA sequences targeting *ANXA1* mRNA (Suppl. Fig. 1). As we have reported previously [31], suppression of *ANXA1* expression led to a baseline increase in paracellular permeability and reduction in TEER. Notably, however, suppression of *ANXA1* expression significantly inhibited the effects of TMAO (40 µM, 24 h) upon both paracellular permeability and TEER (Fig. 3C-D) to a degree that correlated with extent of *ANXA1* suppression across different clones (57/61, 60A and 60B expressing approximately 20, 50 and ∼70 % lower levels of annexin A1, respectively), an effect not seen in cells bearing non-targeting scramble shRNA sequences. The actions of ANXA1 are mediated to a large extent through the G protein-coupled receptor formyl peptide receptor 2 (FPR2) [56]. Hence, we investigated how inclusion of a well-characterised antagonist to this receptor, WRW_4_ (10 µM, 10 min pre-treatment), would affect the functional response to TMAO. Pre-treatment with WRW_4_ was able to significantly attenuate the effects of TMAO treatment on both TEER (Fig. 3E) and paracellular permeability (Fig. 3F), further indicating the role of ANXA1 signalling as the principal mediator of TMAO actions on hCMEC/D3 cells.

### Acute TMAO treatment enhances BBB integrity in vivo

While hCMEC/D3 endothelial cells are a widely used and generally representative model, they cannot reflect all aspects of the multicellular neurovascular unit that underlies BBB function, hence we investigated whether the beneficial effects of TMAO identified *in vitro* translate to an *in vivo* situation. Initial studies revealed that systemic administration of TMAO to wild-type male mice (1.8 mg/kg, i.p.) induced a time-dependent reduction in BBB permeability to the tracer Evans blue (2 % in saline, 100 µl, i.p.), with a significant reduction in dye extravasation to the brain parenchyma being apparent 2 h following TMAO administration, an effect lost at longer time-points (Fig. 4A), presumably due to the relatively short plasma half-life of TMAO *in vivo* [5,57]. To further investigate this effect of TMAO, we employed a simple model of enhanced BBB permeability, namely acute peripheral administration of bacterial LPS [31]. Treatment with LPS (*E. coli* O111:B4, 3 mg/kg, i.p.) significantly enhanced intraparenchymal extravasation of Evans blue within 4 h, an effect significantly attenuated by subsequent treatment with TMAO (1.8 mg/kg, i.p.) 2 h post-LPS (Fig. 4B), further confirming a beneficial action of TMAO at physiological concentrations upon the BBB *in vivo*.

**Fig. 4.**
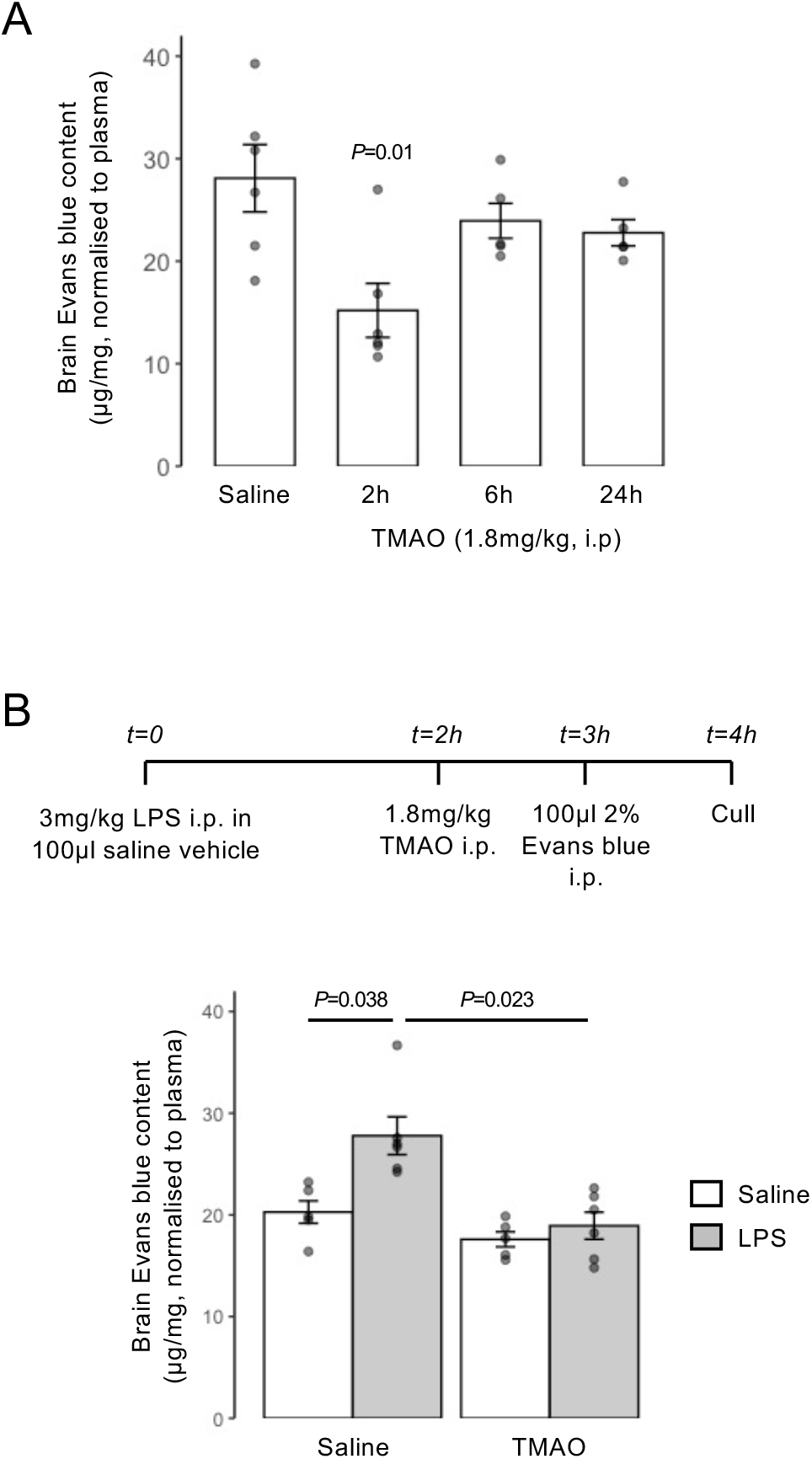
Acute treatment with TMAO promotes BBB integrity *in vivo*. (A) Extravasation of Evans blue dye into brain parenchyma over a 1 h period in 2-month-old male C57Bl/6J mice following i.p. injection of 1.8 mg/kg TMAO for 2 h, 6 h or 24 h *vs.* a saline injected control. Data are normalised to plasma Evans blue content, and are expressed as mean ± s.e.m., *n*=5-6 mice. (B) Extravasation of Evans blue dye into brain parenchyma over a 1 h period in 2-month-old male C57Bl/6J mice following i.p. injection of saline or E. coli O111:B4 LPS (3 mg/kg) with or without subsequent i.p. injection of 1.8 mg/kg TMAO according to the schedule shown. Data are normalised to plasma Evans blue content, and are expressed as mean ± s.e.m., *n*=4-6 mice.

### TMAO treatment rapidly alters brain transcriptional activity

To investigate the wider actions of TMAO upon the brain we performed whole brain RNAseq transcriptomic analysis of wild-type male mice 2 h following TMAO administration (1.8 mg/kg i.p.). We identified 76 significantly differentially expressed genes (P_FDR_<0.1), with expression of 41 upregulated and 35 downregulated (Figure 5A; Supplementary Table 11). KEGG pathway analysis using Enrichr identified a number of significantly regulated murine pathways (Figure 5B), including oxidative phosphorylation, Parkinson’s disease and Alzheimer’s disease. Closer analysis of regulated genes identified several general groupings (Figure 5C), with downregulated genes associated with the mitochondrial respiratory chain (*COX1*, *COX3*, *ATP6*, *ND4L*, *CYTB*, *ND1*, *ND3*, *ND4*, *ND6*) and ribosomal function (*mt-Rnr2*, *mt-Rnr1*, *Rps23rg1*) and upregulated genes associated with cellular or axonal growth (*Nme7*, *B3gat2*, *Fuz*, *Nefm*, *Basp1*, *Mtg1*, *Vps37a*, *Smim1*, *Araf*). Of the 203 BBB-associated human genes previously identified [2], 197 had matches in our mouse brain data set. Here, two genes were identified as significantly differentially expressed at P_FDR_<0.1: reduced *Cpe* (carboxypeptidase E) and increased *App* (amyloid precursor protein) expression (Figure 5D; Supplementary Table 12).

**Fig. 5.**
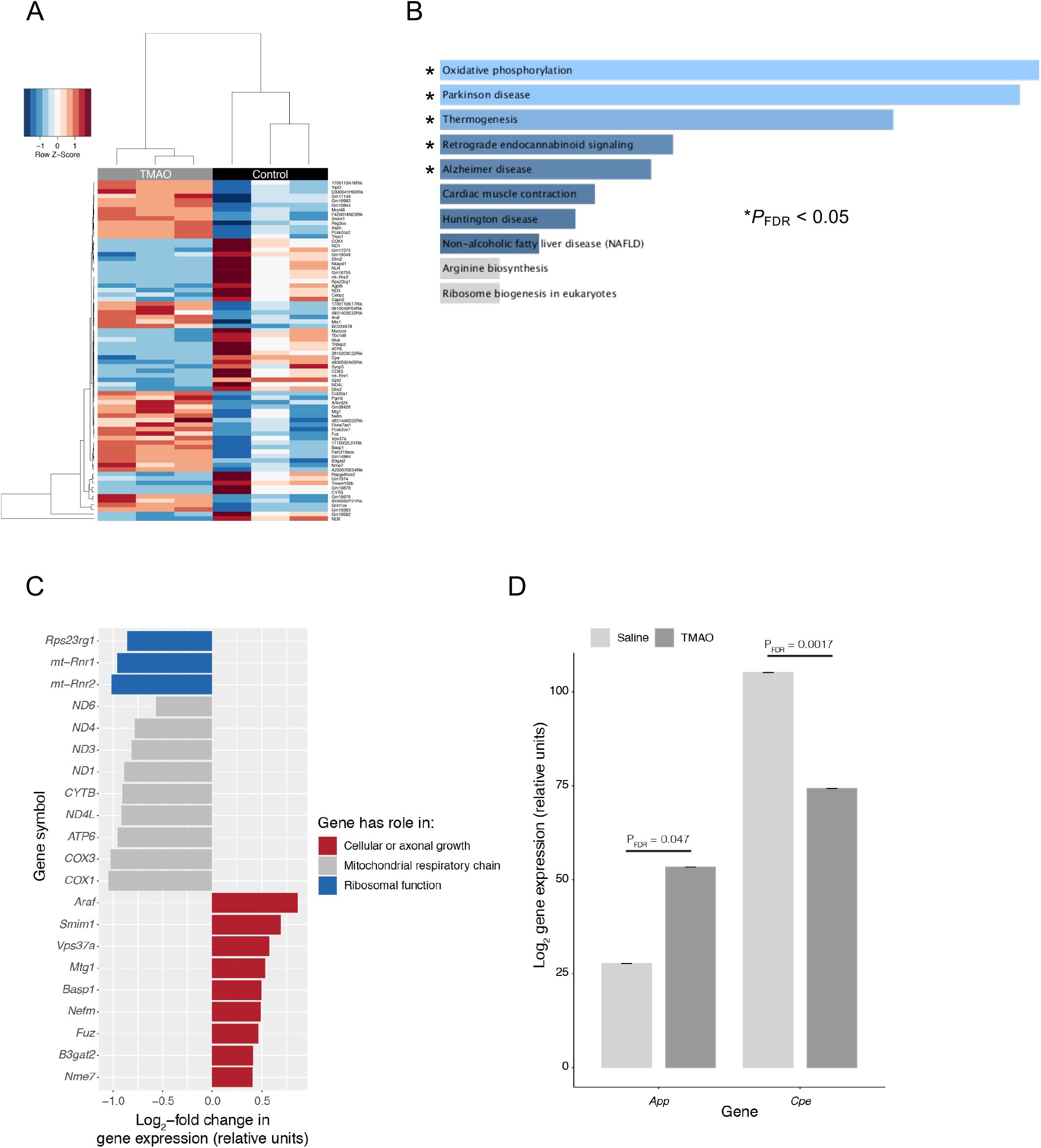
Acute exposure of mice to TMAO significantly alters the whole brain transcriptome. (A) Heatmap showing expression of the 76 genes found to be significantly (P_FDR_<0.1) differentially expressed in the mouse brain after 2 h exposure to 1.8 mg/kg TMAO (*n*=3 per group). Data were scaled by row. (B) Over-representation analysis (Enrichr) showing KEGG pathways associated with the 76 genes. (C) Comparative analysis of significantly differentially expressed genes identified groupings associated with distinct biological functions. (D) Among the 197 BBB-specific genes identified in the data set, only *App* and *Cpe* were significantly (*P*_FDR_<0.1) differentially expressed in the mouse brain after 2 h exposure to TMAO. Data are shown as mean ± s.d, *n*=3 per group. Individual data points are not shown due to the negligible values of the s.d.

### Chronic low-dose TMAO treatment prevents LPS-induced BBB disruption and memory impairment

The fundamental role of the BBB is to protect the brain and preserve its homeostatic environment; damage to BBB integrity is therefore detrimental, and is believed to directly contribute towards cognitive impairment [58]. Having shown TMAO to exert a beneficial effect upon BBB function/integrity in response to acute inflammatory insult, we next examined whether a similar effect held true for chronic conditions, and whether any protection extended to cognition. TMAO was administered to male C57Bl/6J mice through drinking water (0.5 mg/ml) over 2 months, in combination with chronic low-dose LPS administration (0.5 mg/kg/week, i.p.) to model a mild inflammatory stress known to impact cognitive behaviour [43]. There were no differences in volumes of water drunk or, where relevant, final consumption of TMAO between any groups (Table 2). The serum inflammatory cytokines TNFα and IL-1β were both nominally elevated in response to LPS treatment, although not reaching statistical significance, indicating a sub-clinical inflammatory response; TMAO had no effect on TNFα nor IL-1β levels (Suppl. Fig. 4). Notably, animals exposed to LPS exhibited a significant reduction in body weight gain compared to their untreated counterparts, an effect reversed by TMAO treatment (Fig. 6A). Treatment with LPS increased cerebellar FITC extravasation, an effect that was prevented by TMAO treatment, although this did not reach statistical significance on *post hoc* analysis (Fig. 6B). To corroborate these findings, we investigated a second marker of impaired BBB integrity, confocal microscopic detection of brain perivascular IgG deposition. In comparison with sham-treated animals, exposure to LPS caused a significant accumulation of IgG in the perivascular compartment, an effect prevented by TMAO treatment (Fig. 6C).

**Table 2.** Daily consumption of TMAO and water in mice chronically treated with TMAO and/or LPS. Data are mean ± standard deviation.

| Variable | Sham-treated mice | LPS-treated mice | *P* value |
| --- | --- | --- | --- |
| TMAO intake (mg/day/mouse) | 2.79 ± 0.38 | 2.82 ± 0.25 | 0.7 |
| TMAO intake (mg/kg/mouse) | 85.0 ± 11.4 | 88.9 ± 7.9 | 0.14 |
| Water consumption (ml/day/mouse) | 5.57 ± 0.8 | 5.64 ± 0.5 | 0.48 |

**Fig. 6.**
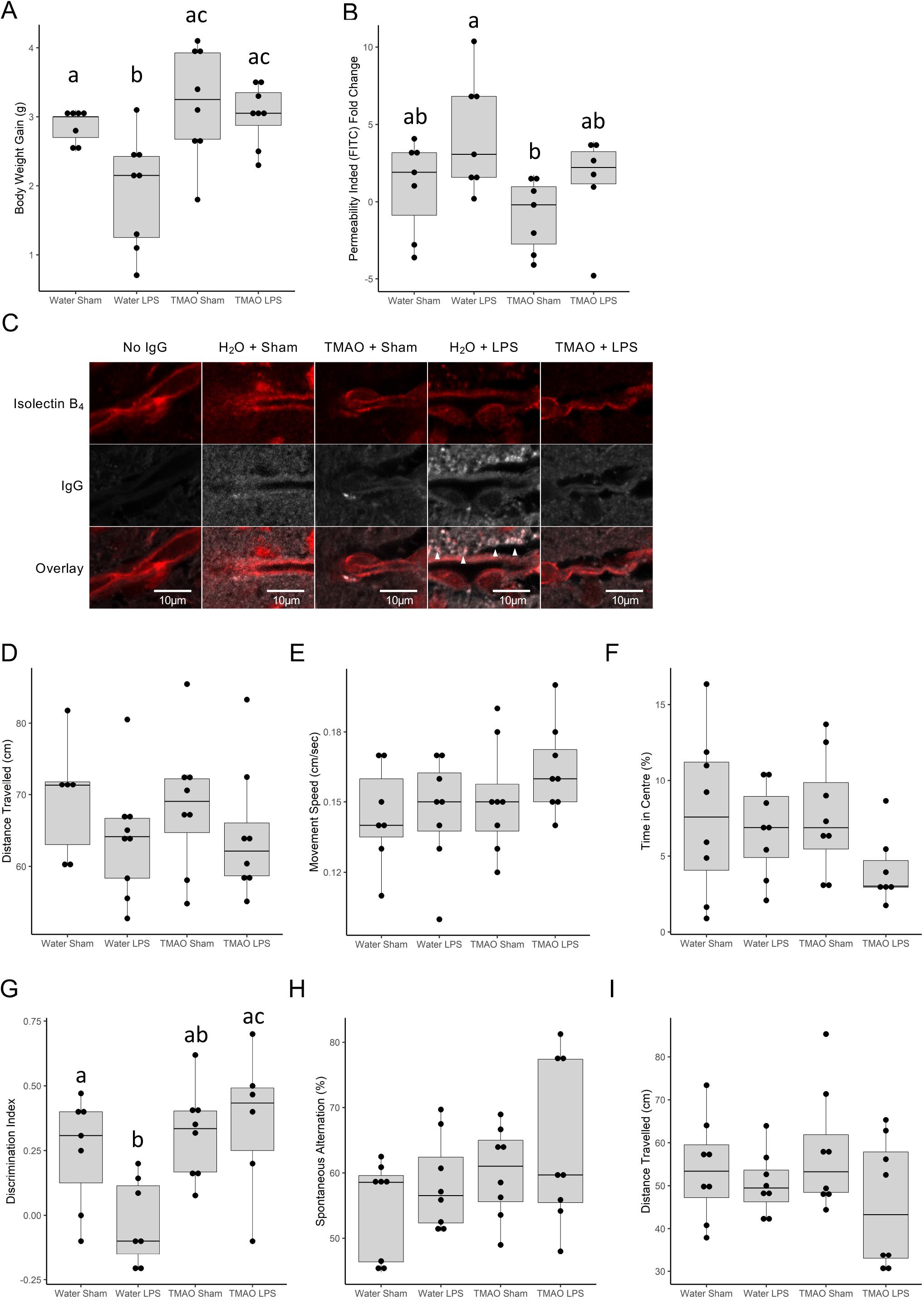
Effect of long-term TMAO exposure on BBB integrity and cognitive function of mice in conjunction with sub-acute inflammatory challenge. (A) Body weight gain in mice treated with TMAO through their drinking water (0.5 mg/ml) over 2 months, combined with a chronic low dose administration of LPS (0.5 mg/kg/week, i.p.). Data are expressed as mean ± s.e.m., *n*=8 mice, columns with different letters are significantly different at *P*<0.05. (B) Cerebellar permeability index to sodium fluorescein 2h following administration in animals previously treated with TMAO through their drinking water (0.5 mg/ml) over 2 months, combined with a chronic low dose administration of LPS (0.5 mg/kg/week, i.p.). Data are expressed as mean ± s.e.m., *n*=8 mice, columns with different letters are significantly different at *P*<0.05. (C) Typical confocal microscopic images of perivascular IgG deposition in male C57Bl/6J mice treated with TMAO through their drinking water (0.5 mg/ml) over 2 months, combined with a chronic low dose administration of LPS (0.5 mg/kg/week, i.p.). *Griffonia simplicifolia* isolectin B_4_ (red) defines endothelial cells, areas of IgG deposition (white) are highlighted by arrow heads. (D) Distance travelled, (E) movement speed and (F) percentage of time in the centre as measured in the OFT in animals previously treated with TMAO through their drinking water (0.5 mg/ml) over 2 months, combined with a chronic low dose administration of LPS (0.5 mg/kg/week, i.p.). Data are expressed as mean ± s.e.m., *n*=8 mice. (G) Novel object discrimination index, calculated as described in Methods, of animals previously treated with TMAO through their drinking water (0.5 mg/ml) over 2 months, combined with a chronic low dose administration of LPS (0.5 mg/kg/week, i.p.). Data are expressed as mean ± s.e.m., *n*=8 mice, columns with different letters are significantly different at *P*<0.05. (H) Percentage of spontaneous alternation and (I) total distance travelled in the Y-maze test for animals previously treated with TMAO through their drinking water (0.5 mg/ml) over 2 months, combined with a chronic low dose administration of LPS (0.5 mg/kg/week, i.p.). Data are expressed as mean ± s.e.m., *n*=8 mice.

The OFT confirmed neither LPS nor TMAO treatment affected motor function, with movement speed and distance travelled comparable across treatment groups (Fig. 6D-E). Similarly, no effect was apparent on the proportion of time animals spent in the centre of the field, suggesting limited effects upon anxiety (Fig. 6F). Working memory, however, determined via NOR indicated a significant reduction in performance in animals exposed to LPS, a behavioural deficit notably prevented in animals co-treated with TMAO (Fig. 6G). In contrast, no effect of either LPS or TMAO treatment was apparent in the Y-maze spontaneous alternation task (Fig. 6H) or in distance travelled during this task (Fig. 6I), indicating no differences in spatial memory.

The brain circuitry thought to underlie spatial and recognition memory functions are known to differ, with greater relative involvement of the hippocampus and entorhinal/perirhinal/retrosplenial cortices, respectively [59,60]. As the underlying cognitive lesion in the NOR task was induced by LPS, we investigated the impact of LPS or TMAO treatment on principal inflammation-responsive CNS cells, astrocytes and microglia, in the entorhinal cortex and hippocampus. Exposure to LPS caused a significant reduction in primary process number for both GFAP+ astrocytes and Iba1+ microglia in the entorhinal cortex, changes that were effectively prevented by TMAO treatment (Fig. 7A-C). Notably, however, no differences were seen in either astrocyte or microglial morphology in the neighbouring hippocampus (Fig. 7E-G); no differences were apparent in either astrocyte or microglial density in either region (Fig. 7D, H).

**Fig. 7.**
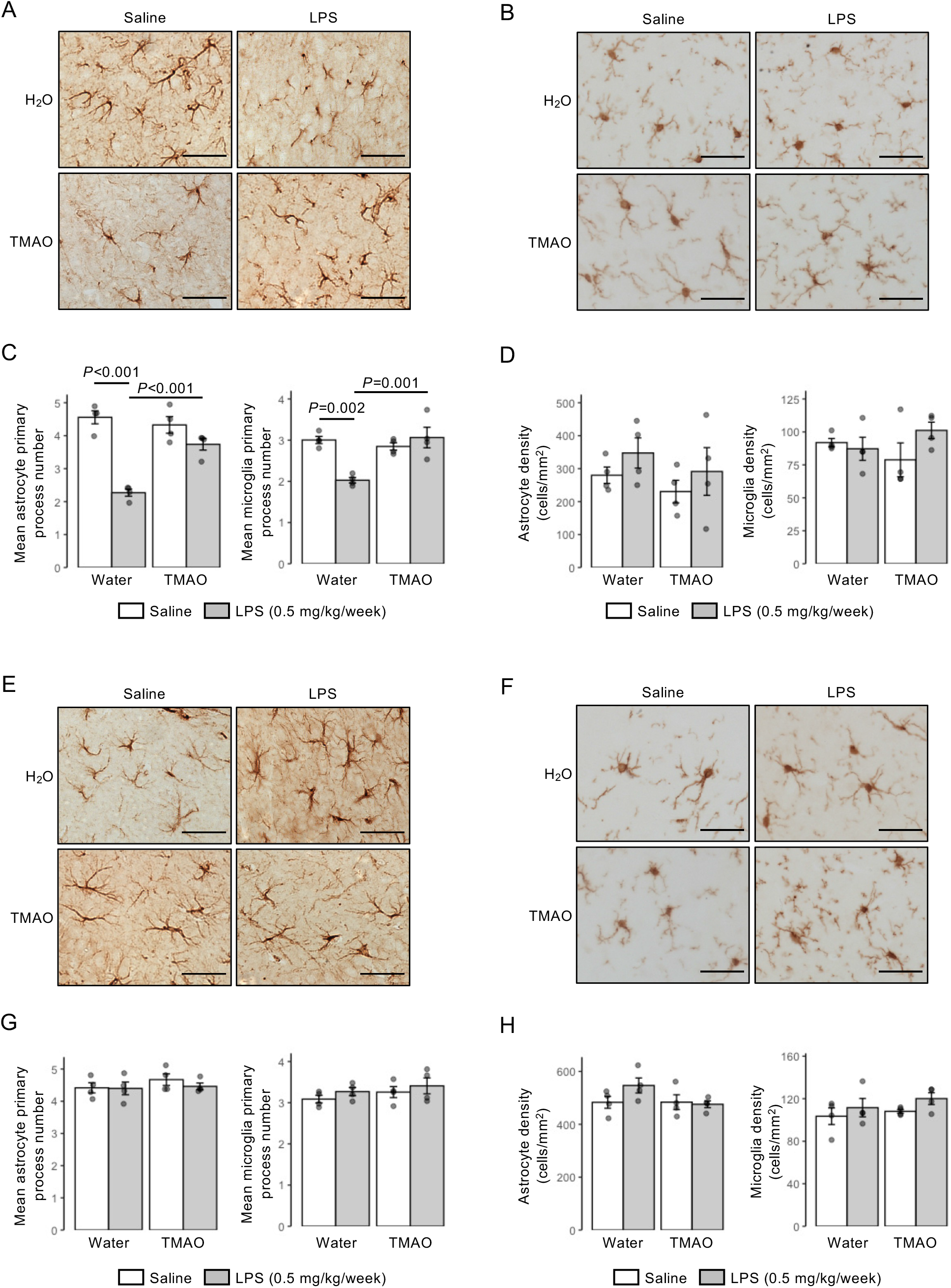
Effects of long-term TMAO exposure upon astrocytes and microglia in the entorhinal cortex and hippocampus of mice in conjunction with sub-acute inflammatory challenge. (A) Typical immunohistochemical staining of GFAP+ astrocytes in the entorhinal cortex of mice previously treated with TMAO through their drinking water (0.5 mg/ml) over 2 months, combined with a chronic low dose administration of LPS (0.5 mg/kg/week, i.p.). Scale bar = 40 µm. (B) Typical immunohistochemical staining of Iba1+ microglia in the entorhinal cortex of mice previously treated with TMAO through their drinking water (0.5 mg/ml) over 2 months, combined with a chronic low dose administration of LPS (0.5 mg/kg/week, i.p.), scale bar = 40 µm. (C) Astrocyte and microglial primary process number and in the entorhinal cortex of mice previously treated with TMAO through their drinking water (0.5 mg/ml) over 2 months, combined with a chronic low dose administration of LPS (0.5 mg/kg/week, i.p.). Data are expressed as mean ± s.e.m., *n*=4 mice. (D) Astrocyte and microglial density in the entorhinal cortex of mice previously treated with TMAO through their drinking water (0.5 mg/ml) over 2 months, combined with a chronic low dose administration of LPS (0.5 mg/kg/week, i.p.). Data are expressed as mean ± s.e.m., *n*=4 mice. (E) Typical immunohistochemical staining of GFAP+ astrocytes in the CA1 region of the hippocampus of mice previously treated with TMAO through their drinking water (0.5 mg/ml) over 2 months, combined with a chronic low dose administration of LPS (0.5 mg/kg/week, i.p.). Scale bar = 40 µm (F) Typical immunohistochemical staining of Iba1+ microglia in the CA1 region of the hippocampus of mice previously treated with TMAO through their drinking water (0.5 mg/ml) over 2 months, combined with a chronic low dose administration of LPS (0.5 mg/kg/week, i.p.), scale bar = 40 µm. (G) Astrocyte and microglial primary process number in the CA1 region of the hippocampus of mice previously treated with TMAO through their drinking water (0.5 mg/ml) over 2 months, combined with a chronic low dose administration of LPS (0.5 mg/kg/week, i.p.). Data are expressed as mean ± s.e.m., *n*=4 mice. (H) Astrocyte and microglial density in the in the CA1 region of the hippocampus of mice previously treated with TMAO through their drinking water (0.5 mg/ml) over 2 months, combined with a chronic low dose administration of LPS (0.5 mg/kg/week, i.p.). Data are expressed as mean ± s.e.m., *n*=4 mice.

## DISCUSSION

The relationship between the BBB and cognitive behaviour is complex and far from being fully understood, but it is clear from both human and animal studies that deficits in barrier integrity can exert a profound and deleterious effect upon memory, language and executive function [61–64]. Indeed, BBB impairment is among the first events to occur in the course of Alzheimer’s disease, and may aggravate the pathological processes that underlie the condition [65]. Strategies to promote BBB function may thus have significant value in helping to protect the brain from progressive neurological diseases such as dementia. In this study we identify novel and distinct roles for the microbiome-associated dietary methylamines TMA and TMAO in regulating BBB function *in vitro* and *in vivo* and provide evidence that the beneficial action of TMAO upon the BBB under inflammatory conditions coincides with similarly positive effects upon glial activity and cognition. These data reinforce the position of the cerebral vasculature as a major target for the gut-brain axis, and extend our knowledge of its interactions with microbial metabolites beyond SCFAs [1,2] to another major class of molecules, the dietary methylamines.

Notably, our data show that while both TMA and TMAO have activity upon the endothelium, there is a marked distinction between their effects despite their close structural similarity. TMA, a volatile organic compound and the direct product of microbial choline, L-carnitine and TMAO metabolism in the upper gut [5], had a deleterious effect upon the endothelium, disrupting cytoskeletal arrangement, inducing signs of metabolic stress and ultimately impairing endothelial barrier integrity. In contrast, TMAO, an inert small molecule largely derived from hepatic FMO3-mediated oxidation of TMA taken up from the gut via the hepatic portal vein [6], promoted cerebral vascular integrity *in vitro* and *in vivo*. These differences suggest that host conversion of TMA (a gas) to TMAO (a stable metabolite) may be an effective detoxification pathway, emphasising the importance of host metabolic pathways in modulating communication in the gut-brain axis, and underlining the importance of using a systems-level approach to understand the interactions between the host and its resident microbiota.

The primary focus of this work was on the effects of TMAO upon the BBB, but this is not necessarily the only CNS target for the methylamine. Our data add to the evidence suggesting that astrocytes [66,67] and microglia [68,69] may respond to TMAO treatment, although it is notable that previous studies have shown pro-activating effects of TMAO at supra-physiological concentrations (>50 μM). An intriguing finding of the current study is the brain region selectivity in the effects of long-term LPS and/or TMAO treatment upon parenchymal glia, with astrocytes and microglia of the entorhinal cortex showing clear LPS-induced, TMAO-sensitive activation, whereas the same cell types in the neighbouring hippocampus appeared resistant to either stimulus. Notably, this closely accords with the involvement of these areas in recognition and spatial memory tasks [59,60], potentially underpinning the cognitive consequences of LPS and TMAO treatment. Determining why this regional discrepancy occurs lies outside the scope of the present study, but it may be relevant that differences have been identified in both neurovascular unit microanatomy [70] and in vascular density [71] between the hippocampus and cortical areas. Ultimately, interpretation of the cell-type-specific responses to TMAO treatment and their interactions with each other, particularly in the context of understanding cognitive implications, will require use of more sensitive analytical techniques such as single-cell transcriptomics, but this remains a fascinating avenue for future study.

Numerous groups have investigated the putative relationship between TMAO and cognition following reports of an association between cerebrospinal fluid TMAO content and Alzheimer’s disease [28], with negative correlations between plasma TMAO content and cognitive function having been identified in both clinical [66,72,73] and experimental [29,68,69] settings. Whether this relationship is truly deterministic remains unclear, however, as the role of the immediate precursor to TMAO – TMA – in cognition and vascular function has largely been overlooked. This omission may be important in light of studies reporting negative correlations between cognitive impairment and serum TMA [74–76] and our own data showing a potent detrimental effect of physiological levels of TMA upon the cerebrovascular endothelium *in vitro*. Given that TMA has also been shown to be detrimental in contexts other than cognitive function [22,23], the contribution that this metabolite plays in disease is evidently in need of closer attention.

Interpreting associations between circulating TMAO and cognition is further complicated by studies indicating that consumption of the TMAO precursors choline and L-carnitine can improve cognitive function [24,25,77], evidence that patients with Parkinson’s disease have lower circulating TMAO than healthy controls [78], and more-recent Mendelian randomisation analysis indicating that serum TMAO and Alzheimer’s disease are not causally related [79]. Given this background, our data indicating that physiologically relevant concentrations of TMAO have positive effects upon both BBB integrity and cognition *in vivo* thus serve as a useful counterweight to population-level correlation studies. Interestingly, a number of previous interventional studies have been performed in mice, suggesting that substantially higher doses of TMAO may have detrimental effects upon learning and memory [68,69,80], although as we and others [81] have identified dose-dependency in the effects of TMAO *in vitro*, it seems plausible that this may reflect a similar phenomenon *in vivo*. The importance of investigating the impact of TMAO under physiologically relevant conditions is further emphasised by a recent study showing TMAO treatment to impair novel object recognition in mice [66], ostensibly an opposite finding to our data achieved with a similar dosing regimen. Importantly, however, mice in this study were maintained on a reduced choline diet, a condition known to alter hepatic metabolism [82]; what impact such changes might have on handling of (TMA and) TMAO by the body is unknown. These discrepancies may be instructive in guarding against incautious extrapolation of TMAO effects from healthy to diseased populations.

Consumption of a diet rich in fish and other seafood, known to provide significant quantities of TMAO [83], associates with a reduced risk of cognitive decline [84,85] and protection against cerebrovascular disease [86]. These effects have in large part been attributed to beneficial actions of the omega-3 polyunsaturated fatty acids [87], although there is little evidence that their direct supplementation improves cognitive function [88] or stroke risk [89]. Here we provide evidence that another component of a seafood-rich diet, TMAO, has protective effects on the cerebral vasculature, astrocyte and microglial function, and upon cognition. Moreover, fish consumption has been associated with reduced inflammatory disease, again attributed primarily to a role for omega-3 fatty acids [90]. While it is too early to definitively claim an anti-inflammatory role for dietary methylamines, particularly given the opposing actions of TMA and TMAO, our data do indicate that broadening the scope of nutritional analyses of seafood-rich diets beyond the omega-3 fatty acids may be worthwhile.

The data we report here indicate a clear reparative effect of TMAO upon BBB integrity following acute inflammatory insult, and further suggest that long-term TMAO treatment may be genuinely protective against prolonged sub-acute inflammatory challenge. Understanding the mechanism(s) underlying these effects is complex, however, as LPS is known to impair BBB function both directly at the endothelium [91] and following systemic cytokine induction [92]. As TMAO acts via induction of ANXA1 release and ANXA1 is known both to enhance BBB integrity [54] and to exert powerful pro-resolving actions at inflammatory foci [93], either/both of these actions of LPS could conceivably be modulated by TMAO treatment. Future studies giving TMAO in advance of BBB challenge may thus be necessary to fully interpret its actions in human clinical settings.

## CONCLUSIONS

Interest in the role played by the gut microbiota in communication through the gut-brain axis has grown dramatically in the last few years, with much attention focused on the mediating actions of microbe-derived metabolites [94]. While a number of studies have shown patterns in microbial metabolite production that associate with different brain functions [95], detailed understanding of the role of individual molecules remains in its infancy, with defined roles characterised for only a subset of the many molecules known to be released by gut microbes. Here we show that the dietary methylamine TMAO can beneficially modulate both BBB integrity and cognitive function *in vivo*, providing direct mechanistic evidence for a positive role of this microbiome-associated metabolite, and reinforce the position of the BBB as an interface in the gut-brain axis. Notably, the positive effects of TMAO that we report stand in contrast to previous work describing deleterious effects of TMAO exposure at high concentrations or under non-physiological conditions [81], emphasising the importance of taking a holistic approach to understanding gut microbiota-host interactions.

## LIST OF ABBREVIATIONS

BBB: blood–brain barrier
DI: discrimination index
GO: gene ontology
KEGG: Kyoto Encyclopedia of Genes and Genomes
LPS: lipopolysaccharide
NOR: novel object recognition
SCFA: short-chain fatty acid
OFT: open field test
SPIA: signalling pathway impact analysis
TEER: transendothelial electrical resistance
TMA: trimethylamine
TMAO: trimethylamine *N*-oxide

## DECLARATIONS

### Ethics approval and consent to participate

Not applicable

### Consent for publication

Not applicable

### Availability of data and materials

Cell line array data have been deposited in ArrayExpress under accession number E-MTAB-6662. Raw murine RNAseq data have been deposited with ArrayExpress under accession number E-MTAB-9869. Supplementary materials associated with the article are available from figshare (https://doi.org/10.6084/m9.figshare.13549334.v1).

### Competing interests

The authors declare that they have no competing interests.

### Funding

This work was funded by Alzheimer’s Research UK Pilot Grant No. ARUK-PPG2016B-6. PREDEASY™ efflux transporter analysis kits were generously provided through the SOLVO Biotechnology Research and Academic Collaborative Transporter Studies (ReACTS) Program. This work used the computing resources of the UK MEDical BIOinformatics partnership – aggregation, integration, visualisation and analysis of large, complex data (UK MED-BIO), which was supported by the Medical Research Council (grant number MR/L01632X/1). SF was supported by Fundación Alfonso Martín Escudero. TS was supported by a bursary from the Imperial College London Undergraduate Research Opportunities Programme.

### Author contributions

LH, DV and SM designed the experiments. SM performed cellular assays and acute *in vivo*/*ex vivo* analyses. TS carried out the initial permeability and TEER assays. KSJ performed glial immunohistochemical analyses. MAA performed IgG extravasation studies. ES produced and provided shRNA treated hCMEC/D3 clones. LH undertook all processing and analyses of transcriptomic data. RCG provided valuable insight and advice throughout the project. DV, MP, IR and MM performed the chronic *in vivo* LPS challenge study and undertook all analyses of behavioural data. SRC, ALC and SF contributed to preliminary animal work. LH, DV and SM wrote the manuscript. All authors read and approved the final version of the manuscript.

## Acknowledgements

Not applicable

## Supplementary Information

**Supplementary Fig. 1.**
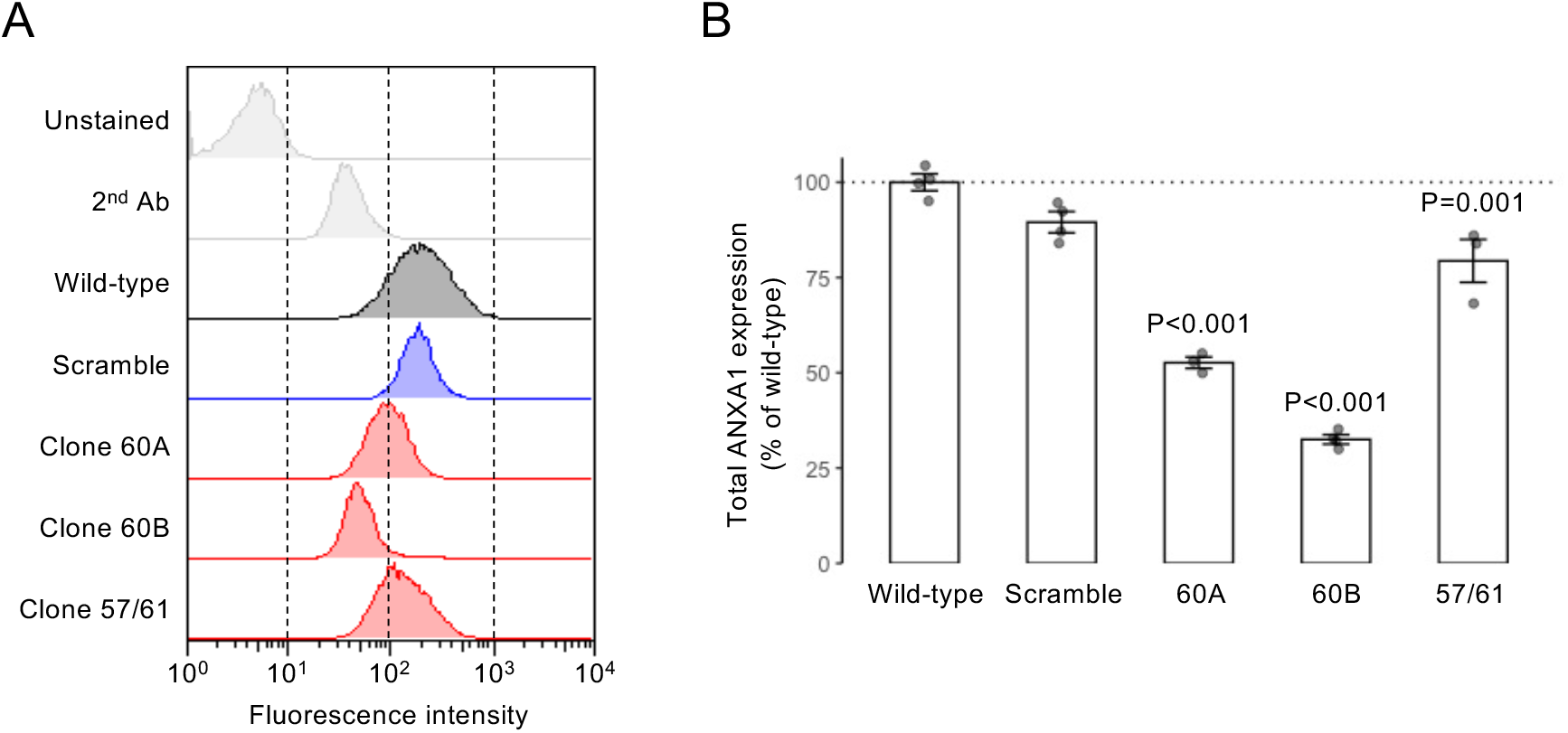
Confirmation of ANXA1 targeting and knock-down in hCMEC/D3 cells stably transfected with appropriate shRNA sequences. (A) Typical flow cytometric profiles of wild-type cells, cells transfected with a scramble shRNA sequence and cells transfected with one of three ANXA1-targeting shRNA sequences, alongside unstained and second antibody controls. (B) Expression of ANXA1 in wild-type hCMEC/D3 cells, cells transfected with a scramble shRNA sequence and cells transfected with one of three ANXA1-targeting shRNA sequences. Data are expressed as mean ± s.e.m., *n*=4 independent experiments.

**Supplementary Fig. 2.**
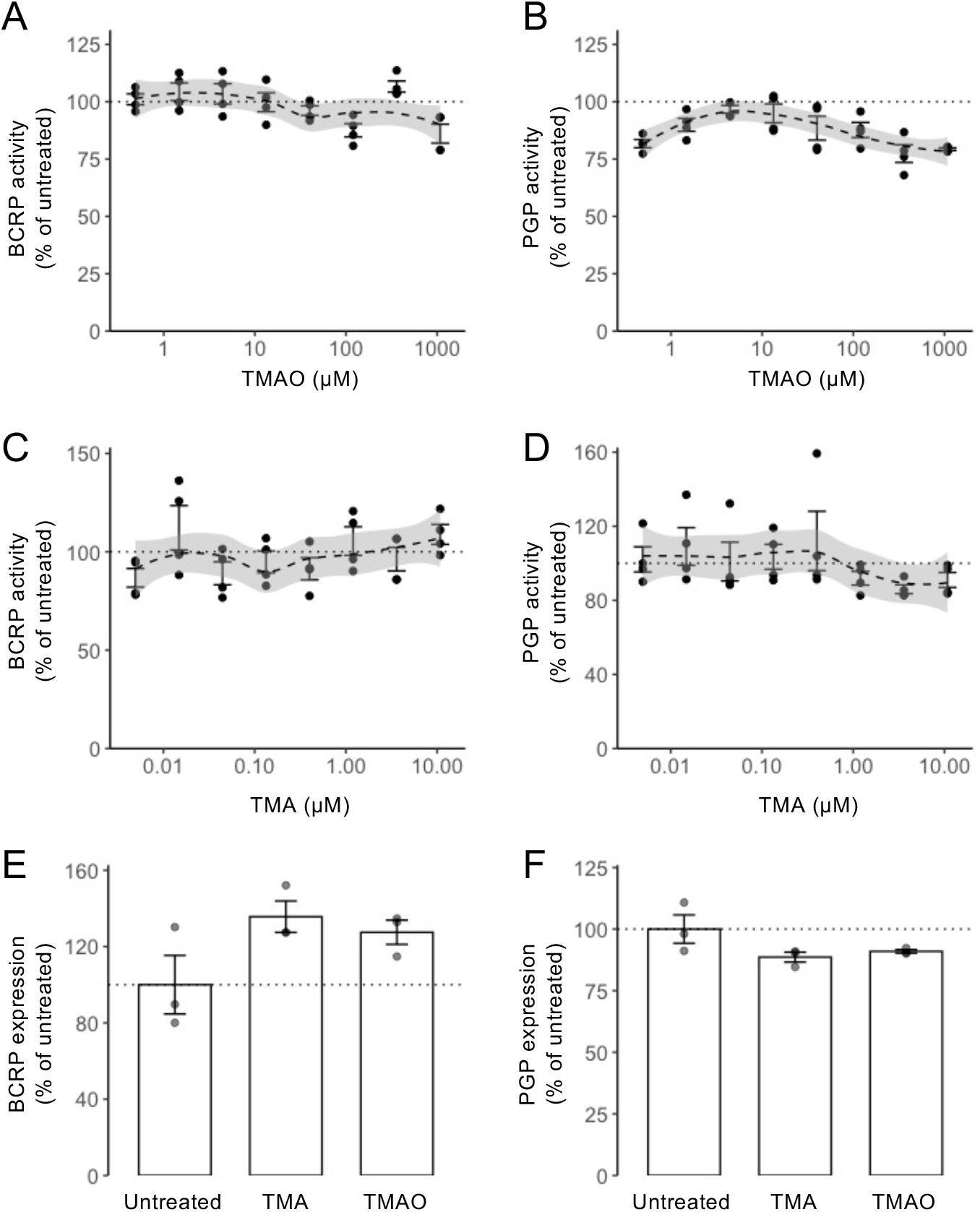
Neither TMAO nor TMA treatment affected the major endothelial efflux transporters BCRP or P-glycoprotein. (A-D) *In vitro* analysis revealed no significant effects of either TMAO or TMA upon BCRP (A, C) or P-glycoprotein (B, D) activity. Dashed lines represent Loess regression fits, with shading representing 95% confidence intervals. (E) hCMEC/D3 cell surface expression of BCRP was unaffected by 24 h exposure to TMA (0.4 µM) or TMAO (40 µM). Data are expressed as mean ± s.e.m., *n*=3 independent experiments. (F) hCMEC/D3 cell surface expression of P-glycoprotein was unaffected by 24 h exposure to TMA (0.4 µM) or TMAO (40 µM). Data are expressed as mean ± s.e.m., *n*=3 independent experiments.

**Supplementary Fig. 3.**
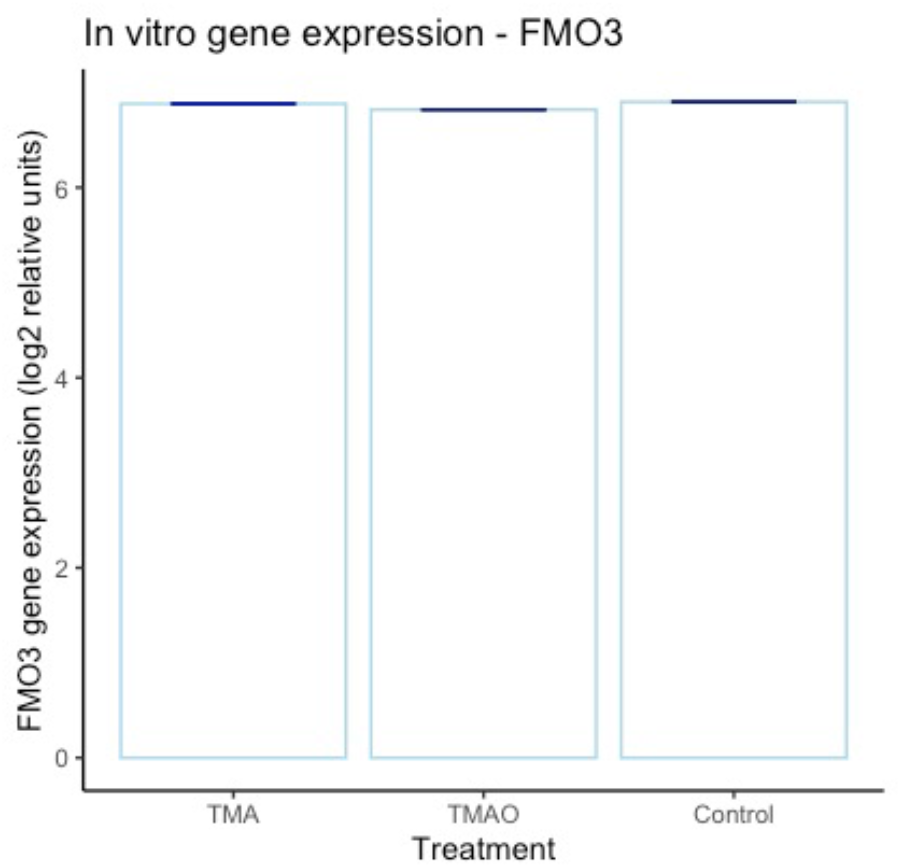
*In vitro* (hCMEC/D3 cell) *FMO3* gene expression is unaffected by exposure to TMA (0.4 µM) or TMAO (40 µM). Dark blue lines represent the standard deviation (+/-). There was no statistically significant difference between TMA and the control, nor between TMAO and the control (Supplementary Tables 2 and 3). Individual data points are not shown due to the negligible values of the standard deviations.

**Supplementary Fig. 4.**
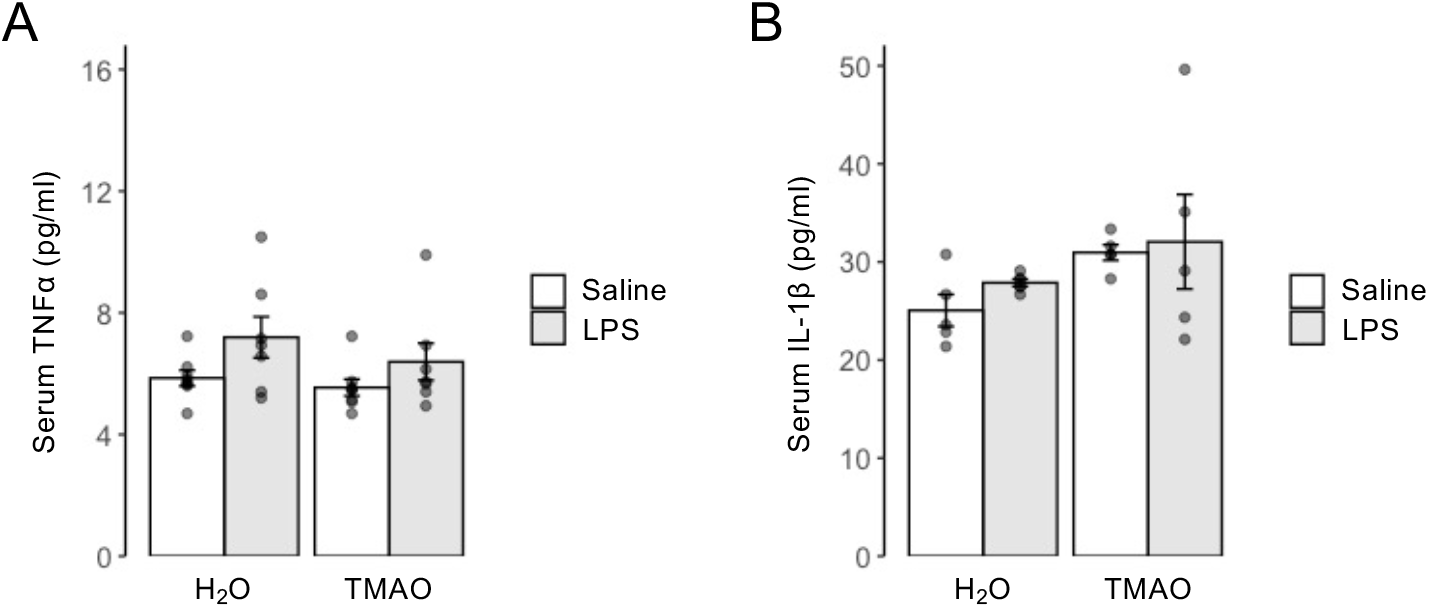
Neither chronic low-dose LPS nor TMAO treatment significantly increases serum inflammatory cytokines. (A) Serum TNFα concentrations in male C57Bl/6J mice treated with TMAO through their drinking water (0.5 mg/ml) over 2 months, combined with a chronic low dose administration of LPS (0.5 mg/kg/week, i.p.). Data are expressed as mean ± s.e.m., *n*=7-8 mice. (B) Serum IL-1β concentrations in male C57Bl/6J mice treated with TMAO through their drinking water (0.5 mg/ml) over 2 months, combined with a chronic low dose administration of LPS (0.5 mg/kg/week, i.p.). Data are expressed as mean ± s.e.m., *n*=7-8 mice.

The supplementary tables listed below are available from figshare (https://doi.org/10.6084/m9.figshare.13549334.v1).

**Supplementary Table 1.** Normalized microarray data for hCMEC/D3 treated or not with TMA (0.4 µM) or TMAO (40 µM)

**Supplementary Table 2.** Differential gene expression for control compared with TMA treatment, as assessed using LIMMA

**Supplementary Table 3.** Differential gene expression for control compared with TMAO treatment, as assessed using LIMMA

**Supplementary Table 4.** KEGG-based SPIA of genes whose expression was significantly affected by TMAO

**Supplementary Table 5.** GO (biological process) analysis of TMA-up-regulated genes as assessed using Enrichr

**Supplementary Table 6.** GO (biological process) analysis of TMA-down-regulated genes as assessed using Enrichr

**Supplementary Table 7.** GO (biological process) analysis of TMAO-up-regulated genes as assessed using Enrichr

**Supplementary Table 8.** GO (biological process) analysis of TMAO-down-regulated genes as assessed using Enrichr

**Supplementary Table 9.** Differential gene expression of BBB-associated genes (*in vitro* TMA treatment)

**Supplementary Table 10.** Differential gene expression of BBB-associated genes (*in vitro* TMAO treatment)

**Supplementary Table 11.** Differential gene expression in mouse brain for control compared with TMAO treatment, as assessed using DESeq2

**Supplementary Table 12.** Differential expression of BBB-associated genes in mouse brain for control compared with TMAO treatment, as assessed using DESeq2

## Notes

https://doi.org/10.6084/m9.figshare.13549334.v1

### Competing Interest Statement

The authors have declared no competing interest.

